# Expression, Purification and Preliminary Analysis of REcTO4, an Engineered Chimeric PIN-Domain Endoribonuclease

**DOI:** 10.64898/2026.11.20.768124

**Authors:** Bianca Braggiotti, Paolo Cocomazzi, Silvia Ferrara, Lucia Corcione, Luca Muzio, Eloise Mastrangelo, Mario Milani

## Abstract

REcTO4 (R4) is an engineered chimeric protein that combines the PIN endoribonuclease domain of Regnase-1 with the tandem zinc-finger (TZF) RNA-binding domain of tristetraprolin (TTP), designed to promote the targeted degradation of pro-inflammatory mRNAs. Recombinant production and purification of this bifunctional chimeric protein in soluble form presents substantial challenges, including the need to accommodate two structurally distinct functional domains that must fold in the bacterial cytoplasm, as well as the interaction of the TZF domain with endogenous RNA. Here, we describe the systematic expression, screening, optimization of purification procedures, and biochemical/biophysical characterization of recombinant GST-tagged R4. Among the five expression constructs designed, only the N-terminally GST-tagged construct yielded an appreciable amount of soluble, catalytically active protein in *Escherichia coli*. An optimized two-step purification protocol, combining GST affinity and size-exclusion chromatography (SEC), yielded GST-R4 with a purity exceeding 95%. GST-R4 also co-purifies with endogenous bacterial RNA, forming high-molecular-weight complexes with apparent molecular mass of approximately 570 kDa by SEC. Endoribonuclease assay performed after affinity purification showed detectable RNA-cleavage activity. Preliminary cryo-EM micrographs reveal heterogeneous particles of approximately 10 nm, together with filamentous features consistent with assemblies of RNA-protein complexes, but intrinsic sample heterogeneity currently prevents 2D/3D reconstruction. This methodology reported here establishes an expression and purification workflow for GST-R4 and provides a foundation for structural and functional studies of this novel chimeric enzyme class engineered for pharmacological application.

**Highlights:**

- Only the GST-tagged R4 construct yields soluble protein in *E. coli*.
- Two-step purification gives >95% pure GST-R4 in a single working day.
- GST-R4 co-purifies with endogenous bacterial RNA with an estimated size of ~200 nt.
- GST-R4 shows endoribonuclease activity *in vitro*.
- Preliminary cryo-EM micrographs reveal heterogeneous particles of approximately ~10 nm.

## 1. Introduction

Post-transcriptional regulation of pro-inflammatory gene expression plays a central role in controlling the magnitude and resolution of innate immune responses (Uehata et al., 2013; Makita et al., 2021). Two key RNA-binding proteins involved in this regulatory network are Regnase-1 (MCPIP1/ZC3H12A) and tristetraprolin (TTP/ZFP36), which promote the degradation of pro-inflammatory mRNAs through distinct molecular mechanisms (Takeuchi et al., 2018). TTP recognizes AU-rich elements (AREs) within the 3′ untranslated regions (3′UTRs) of target transcripts through its tandem zinc-finger (TZF) domain, promoting mRNA deadenylation and subsequent decay through the recruitment of the CCR4-NOT complex (Lai et al., 1999; Makita et al., 2021). In contrast, Regnase-1 acts as an endoribonuclease through its N-terminal PIN-like domain, recognizing specific stem-loop structures within the 3′UTRs of target mRNAs promoting their direct cleavage. Although these proteins regulate partially overlapping inflammatory pathways and can affect the expression of common inflammatory mediators, they exhibit distinct RNA-recognition properties and degradation mechanisms (Takeuchi et al., 2018). Furthermore, their activity is dynamically regulated by inflammatory signaling pathways and post-translational modifications, allowing the fine-tuning of inflammatory gene expression throughout the course of the immune response (Mahtani et al., 2001; Akaki et al., 2021). REcTO (REgnase1–TristetraprOlin) chimeras are a class of rationally engineered proteins designed to combine the RNA-binding specificity of the TTP TZF domain with the endoribonuclease activity of the Regnase-1 PIN domain in a single polypeptide chain. The design rationale is to co-localize sequence-selective RNA recognition with endonucleolytic cleavage activity, thereby enabling targeted destabilization of pro-inflammatory mRNA substrates. Among the chimeric variants explored, REcTO4 (R4) emerged as the lead candidate and was selected for further biochemical and structural characterization; its complete functional characterization will be reported separately.

The chimeric architecture of R4 was developed through structural modelling and iterative construct design to preserve the spatial organization of the PIN domain while accommodating the adjacent TZF region. However, the recombinant production of this multi-domain chimeric protein in a bacterial expression system presents intrinsic challenges that are qualitatively different from those encountered with single-domain targets. The two functional domains of R4 (the PIN endoribonuclease and the TZF RNA-binding domain) are structurally distinct and are derived from proteins with very different physicochemical properties and cellular localization signals. Ensuring the concurrent, correct folding of both domains in the bacterial cytoplasm, while preventing aggregation and preserving the zinc-dependent TZF fold, required extensive construct and expression optimization. Here we report the systematic screening of five expression constructs and multiple purification strategies, and an optimized two-step purification protocol for the GST-R4 fusion protein. We further describe the biochemical and biophysical properties of the purified complex, assess the endoribonuclease activity of the affinity-purified preparation as a functional quality-control step, and report the results of extensive crystallization trials and preliminary cryo-electron microscopy experiments, which provide an initial assessment of the suitability of the purified material for structural studies.

## 2. Materials and Methods

### 2.1 Computational modelling and linker design

The three-dimensional architecture of R4 (murine form) was designed starting from the available crystal structures of the MCPIP1 N-terminal domain (PDB entries 3V33 and 5H9W) as a model for the Regnase-1 PIN domain, together with the NMR structure of the tandem zinc-finger (TZF) domain of TIS11d bound to RNA (PDB entry 1RGO) as a model for the TTP TZF domain (Hudson et al., 2004; Mino et al., 2015). Molecular models were built to evaluate the spatial compatibility of the two domains and to identify candidate junction sequences. The critical design principle was derived from the wild-type Regnase-1 architecture, in which the PIN domain is connected to its intrinsic zinc-finger motif by a short linker of five residues. This compact geometry was adopted as a template for the chimeric junction. The final R4 design employs a direct connection of Regnase-1 Pro298 to TTP Arg95, exploiting the conformational plasticity of the short TSS tripeptide linker at the N-terminus of the TZF domain.

### 2.2 Molecular dynamics simulation

The simulation was performed with the program GROMACS (Pronk et al., 2013) with the AMBER94 force field (Lindorff-Larsen et al., 2010). The initial model, in the presence of one Mg^+2^ and two Zn^+2^, was solvated with 17,945 water molecules, neutralized with 36 Na^+^ and 20 Cl^-^ ions; after energy minimization and system equilibration (NVT) the productive run (NPT) lasted for 180 ns, with 2 fs time step.

### 2.3 Construct design and cloning

To identify a soluble, functional expression system for R4, four distinct constructs were designed and cloned: 1. pET-21a(+)-R4 (43.7 kDa) encodes R4 with an N-terminal His-tag and a PreScission protease site; 2. pGEX-R4 (68.4 kDa) carries an N-terminal GST fusion partner followed by a PreScission site; 3. pGEX-R4-ED is identical to pGEX-R4 with two additional C-terminal residues (Glu-Asp) intended to add stability to the second zinc finger; 4. an aldolase-tagged variant (Aldo-R4) was also explored, employing a custom bacterial aldolase fusion tag reported by Ferrara et al. (Ferrara et al., 2025) as a tool to favor both crystallization and cryo-EM experiments. The plasmids were purchased from ProteoGenix. The complete amino acid sequence of pGEX-R4 (GST fusion, PreScission site, PIN domain and TZF domain) is reported in Figure 1; this construct also contains an internal thrombin site (LVPR↓GS (Xu et al., 2012)) within the Regnase-1 sequence.

**Figure 1.**
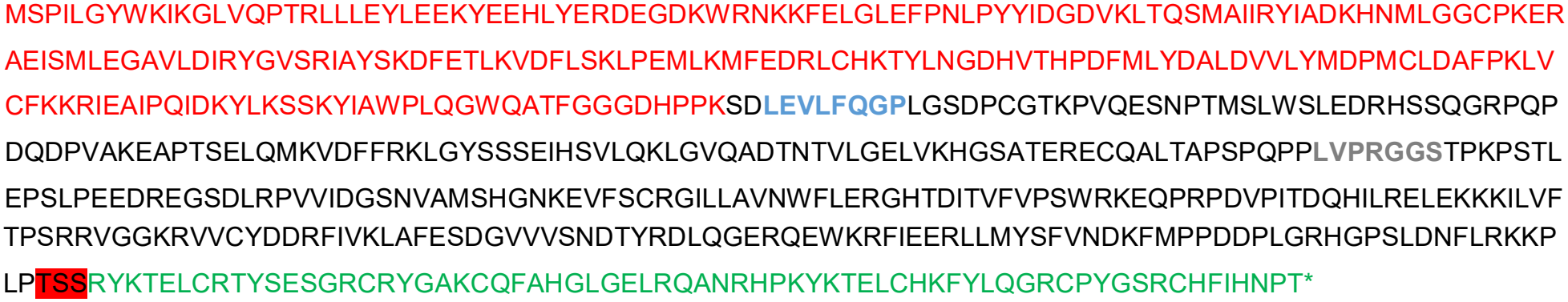
GST-R4 sequence: GST (red), PreScission protease cleavage site (bold blue), thrombin cleavage site within the R4 construct (bold grey), Regnase-1 PIN domain (black), linker (highlighted in red), TTP zinc fingers (green).

### 2.4 Expression screening

Expression trials were carried out in three *E. coli* strains (BL21(DE3), Rosetta and Shuffle), across a range of temperatures (16, 25 and 37°C) and induction times (3 h, 6 h, and overnight) after addition of 0.4 mM IPTG to LB medium cultures. Solubility was assessed by SDS-PAGE of total lysate and soluble fraction after centrifugation at 20,000 × g for 30 min. Among the constructs tested, only pGEX-R4 yielded appreciable amounts of soluble protein, using E. coli BL21(DE3) cells cultured at 16°C overnight following IPTG induction.

### 2.5 Protein Purification: Optimization and Refinement

All purification steps were performed at 4°C on the same day to minimize protein aggregation and preserve protein stability. Cells from 1–2 L cultures were resuspended in lysis buffer (25 mM HEPES pH 7.5, 500 mM NaCl, 1 mM DTT) supplemented with DNase I and protease inhibitors (cOmplete, Roche), lysozyme, benzonase and lysed by sonication. The clarified lysate (20,000 × g, 45 min) was applied to a glutathione-Sepharose affinity column (GSTrap HP, Cytiva). After an extensive wash of approximately 30 column volumes in lysis buffer to maximize purity and remove co-purifying contaminants, the bound protein was eluted with 50 mM HEPES pH 7.5, 200 mM NaCl, 20 mM reduced glutathione, and 1 mM DTT. Peak fractions (estimated purity of approximately 70% by SDS-PAGE) were pooled and immediately subjected to size-exclusion chromatography (SEC) on a HiLoad 16/600 Superdex 200 column (Cytiva) pre-equilibrated with the final storage buffer: 25 mM HEPES pH 7.5, 200 mM NaCl, 1 mM DTT, and 50 μM ZnSO□. Zinc supplementation was included to stabilize the C-terminal zinc-finger motifs. Fractions showing >95% purity by SDS-PAGE were pooled and used for all downstream analyses.

Several buffer conditions were additionally tested in an attempt to improve protein stability. These included: (i) 20 mM Tris, pH 8.0, 300 mM NaCl, under which the protein showed a tendency to degrade; (ii) 50 mM HEPES, pH 7.0, 500 mM NaCl, which promoted protein aggregation and precipitation; (iii) 50 mM MES, pH 6.5, 200 mM NaCl, 25 μM ZnSO, 1 mM DTT, which also resulted in protein aggregation and precipitation; and (iv) 50 mM MES, pH 6.5, 200 mM NaCl, 50 mM glutamate, and 50 mM arginine, under which the protein aggregated and could not be cleaved by either PreScission protease or thrombin.

GST-tag removal was attempted using PreScission protease and thrombin under different conditions, both following affinity purification and after SEC. However, cleavage was unsuccessful or resulted in protein precipitation.

### 2.6 Biochemical and biophysical characterization

Protein concentration was determined by UV absorbance at 280 nm using the theoretical extinction coefficient calculated from amino acid composition (79760; https://web.expasy.org/cgi-bin/protparam/protparam). Nucleic acid co-purification was quantified from the A260/A280 ratio and confirmed by agarose gel electrophoresis. The nature of the co-purified nucleic acid was assessed by parallel treatment of aliquots with RNase T1 or RNase-free DNase I (RQ1 DNase) at 37°C for 30 min, followed by 0.8% agarose gel electrophoresis. The apparent molecular weight of the GST-R4/RNA complex was estimated from SEC elution volume (Ve □ 49.9 ml) against a standard calibration curve. Thermal stability assays were conducted in a StepOne Real Time PCR System (Applied biosystem Thermo Fisher Scientific), at two ionic strengths (200 mM and 1 M NaCl) to probe the RNA-stabilization effect. Samples were heated from 20 to 95 °C (0.2 °C/min), and fluorescence intensity was measured using the Sypro Orange excitation/emission ranges of 470–505/540–700 nm, respectively.

### 2.7 Monitoring of endoribonuclease activity during purification

The endoribonuclease activity of GST-R4 was monitored after affinity chromatography using a fluorescently labelled RNA substrate derived from the 3′ untranslated regions of human IL-6 mRNA. The cleavage assay was designed with reference to previously described in vitro assays for Regnase-1, with modifications (Yokogawa et al., 2016). The substrate consisted of a 5′-Cy5-labelled *IL-6* RNA (5′-UGUUGUUCUCUACGAAGAACUGACA-3′, 25 nt). Cleavage reactions were performed in a final volume of 50 μL containing 20 pmol of RNA substrate (0.4 μM) and 7.5 μg of affinity-purified GST-R4, in cleavage buffer consisting of 20 mM Tris-HCl (pH 7.5), 200 mM NaCl, 1 mM DTT and 25 μM ZnSO□. Reactions were incubated at 37°C, and aliquots were collected at 0, 5, 15, 30, 60 and 120 min. At each time point, 5 μL aliquots were mixed with 5 μL of denaturing loading buffer containing 5 mM EDTA, 0.025% xylene cyanol and 0.025% bromophenol blue in formamide, and heated at 95°C for 5 min. Samples were subsequently analyzed by electrophoresis on 12.8% polyacrylamide (19:1) denaturing TBE-urea gels containing 7 M urea. Cy5-labelled RNA substrate and cleavage products were visualized using a ChemiDoc imaging system.

### 2.8 Crystallization trials

Crystallization experiments were set up with an ORYX-4 crystallization robot (Douglas Instruments Ltd) in sitting-drop and microbatch formats. A total of ~1000 conditions were screened at protein concentrations of 2–6 mg/ml, using commercial sparse-matrix and grid screens (Hampton Research, Molecular Dimensions, Qiagen). To reduce sample heterogeneity from co-purified RNA, additional trials were performed with sub-stoichiometric amounts of RNase A added to the crystallization drop. Synchrotron data were collected at the ESRF beamline ID30A-3 (Grenoble, France) for selected crystal candidates.

### 2.9 Cryo-EM sample preparation and data collection

GST-R4 samples at concentrations of 0.5 and 1.0 mg/ml were applied to glow-discharged Quantifoil R 1.2/1.3 holey-carbon grids (Cu, 300 mesh), blotted and plunge-frozen in liquid ethane using a Vitrobot Mark IV (Thermo Fisher Scientific). Four grids were prepared under varying blotting conditions. Cryo-EM micrographs were collected on a FEI Talos Arctica 200 kV transmission electron microscope at liquid-nitrogen temperature at the cryo-EM facility of the Department of Biosciences, Università degli Studi di Milano (https://sites.unimi.it/biolstru/em.html).

## 3. Results

### 3.1 Design of the R4 chimeric junction

Structural modelling selected the tripeptide junction TSS between Regnase-1 Pro298 and Arg95 TTP (…KPLP – TSS – RYKT…) as the minimal geometry consistent with the native sequence of the PIN domain fold with its proper Zn-finger domain. The inherent conformational plasticity of the TSS tripeptide at the N-terminus of the TTP TZF domain was predicted to accommodate the chimeric architecture in a fold analogous to the native Regnase-1 zinc-finger motif. This R4 construct was taken forward for expression and purification.

### 3.2 Molecular dynamics simulation on the R4 construct

The 3D model of R4 structure, starting after the thrombin cleavage site (GSTP…) was prepared using the PDBs 5H9W and 3V33 for the regnase1 pin domain and 1RGO for the two TTP Zn-fingers. The initial structure after energy minimization and equilibration was simulated for 180 ns to check for consistency of the chosen linker. After ~75 ns the r.m.s.d. respect to the initial conformation remains stable at ~2 A. The linker allows R4 to reach a stable conformation, more compact respect to the initial configuration (Figure 2).

**Figure 2.**
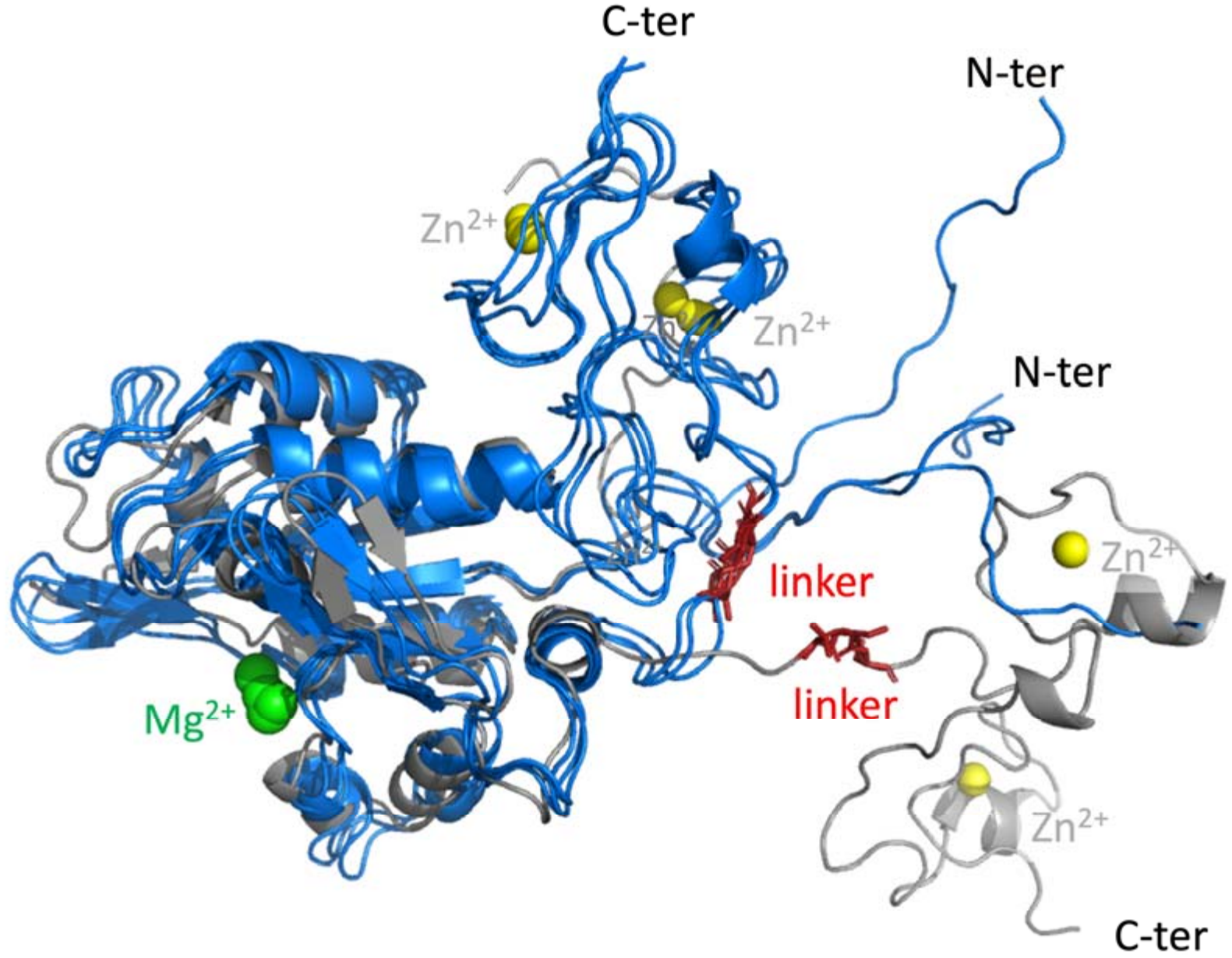
MD simulation: initial model in gray and final conformations at 170, 175 and 180 ns. Mg and Zn are represented as green and yellow spheres, respectively; the TSS linker is depicted as red sticks. Figure prepared using PyMol.

### 3.3 Construct screening and expression optimization

Of the five constructs tested and described in Section 2.2, only pGEX-R4 yielded appreciable amounts of soluble protein. All other constructs [pET-21a(+)-R4 (His-tagged), pGEX-R4-ED (two-residue C-terminal extension) and the Aldo-tagged variant] accumulated in inclusion bodies or failed to express detectably under all conditions explored. The optimal expression conditions for pGEX-R4 were *E. coli* BL21(DE3) grown at 16°C overnight following IPTG induction. Expression at higher temperatures (25°C or 37°C) or in alternative strains (Rosetta, SHuffle) gave predominantly insoluble protein. The critical role of the GST tag in solubilizing R4 indicates that the fusion partner acts as a chaperone-like solubility enhancer for the chimeric PIN/TZF architecture, in addition to providing the affinity handle for purification.

### 3.4 Optimized purification of R4

Extensive buffer screening (described in Section 2.4) revealed that GST-R4 is markedly prone to aggregation, precipitation, and proteolytic degradation. Conditions tested included systematic variation of pH, ionic strength, divalent cations (Ca^2^□, Mg^2^□, Zn^2^□), solubility enhancers (arginine, glutamate) and detergents, as well as the addition of protease inhibitors at every purification step. The two-step protocol described in Section 2.4 (GST-affinity chromatography with an extended wash, followed immediately by SEC) gave the most reproducible results, with all steps completed in a single working day to minimize degradation (Figure 3A-D). SDS-PAGE analysis of SEC peak fractions confirmed purity >95% (Figure 3D). The SEC elution profile shows the protein migrating at a Ve of 49.9 (Figure 3C) corresponding to an apparent molecular weight of 570 kDa, far above the theoretical mass of monomeric GST-R4 (68.4 kDa), indicating the formation of high-order oligomeric assemblies. Starting from 1-2 L bacterial cultures, this protocol consistently delivers material suitable for biochemical assays and structural characterization trials. GST-tag removal using PreScission protease or thrombin was unsuccessful under all conditions tested, preventing the recovery of stable tag-free protein likely due to the multimeric assembly.

**Figure 3.**
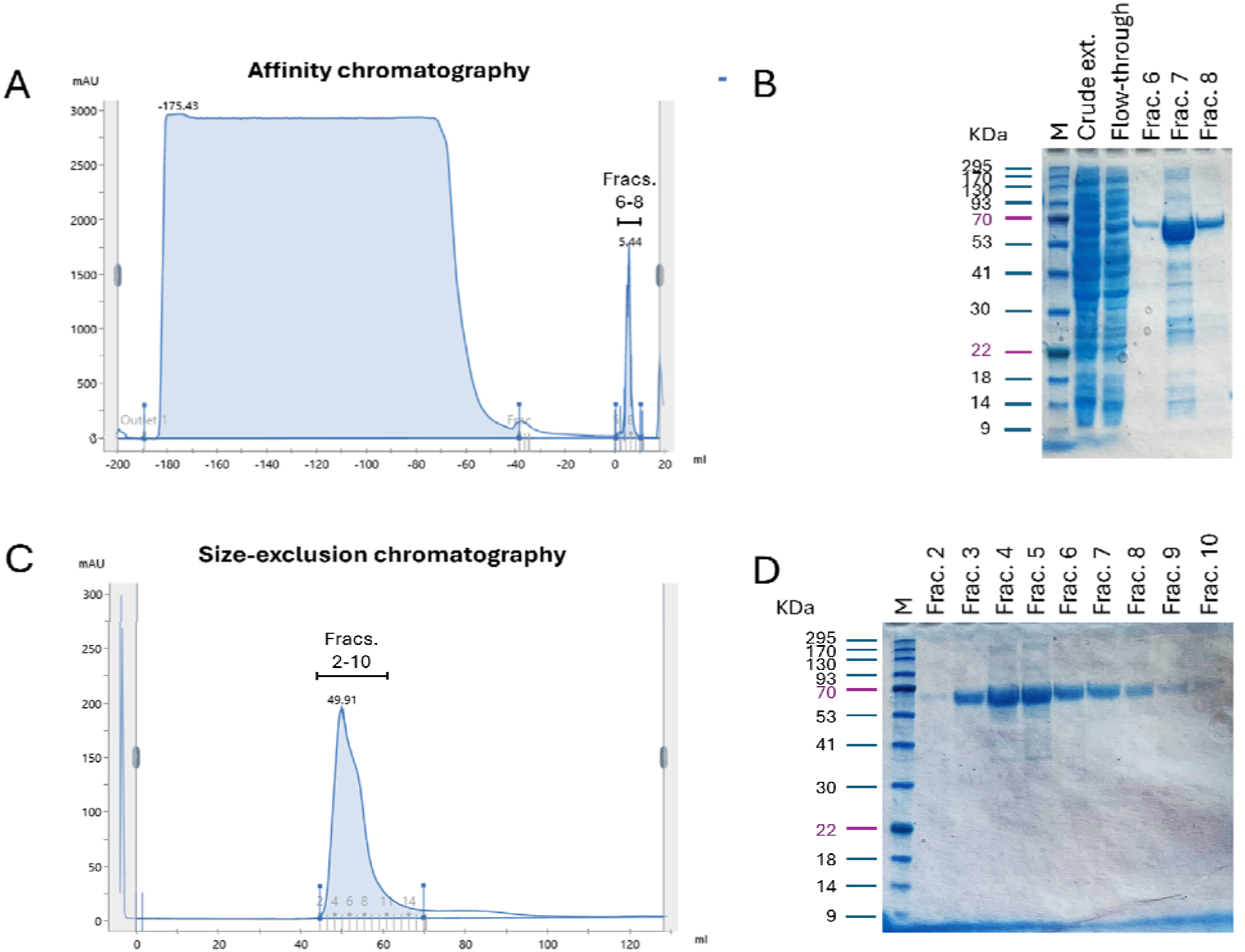
Purification of the GST-R4 construct using the optimized two-step chromatography protocol. (A) Representative affinity chromatography profile obtained using a GSTrap HP column. Fractions 6–8 collected from the elution peak are indicated. (B) Coomassie Brilliant Blue-stained SDS-PAGE analysis of the affinity chromatography samples: molecular-weight marker (M), crude extract, flow-through, and elution fractions 6, 7 and 8. (C) Representative size-exclusion chromatography (SEC) profile obtained using a Superdex 16/600 column. Fractions 2–10 collected across the elution peak are indicated. (D) Coomassie Brilliant Blue-stained SDS-PAGE analysis of the SEC fractions (fractions 2–10). Molecular-weight markers are indicated in kDa.

### 3.5 GST-R4 co-purifies with endogenous bacterial RNA

Absorption spectroscopy of purified GST-R4 preparations consistently showed A260/A280 ratios of 1.6-1.8, suggesting the presence of co-purifying nucleic acids. This signal persisted despite DNase I treatment of the cell lysate during preparation. Samples collected from different fractions during GST-affinity chromatography and subsequent size-exclusion chromatography (SEC) were analyzed by agarose gel electrophoresis (Figure 4A). Nucleic acid bands were detected in samples from both purification steps, indicating that nucleic acid material remained associated with the GST-R4 preparations throughout the purification process.

**Figure 4.**
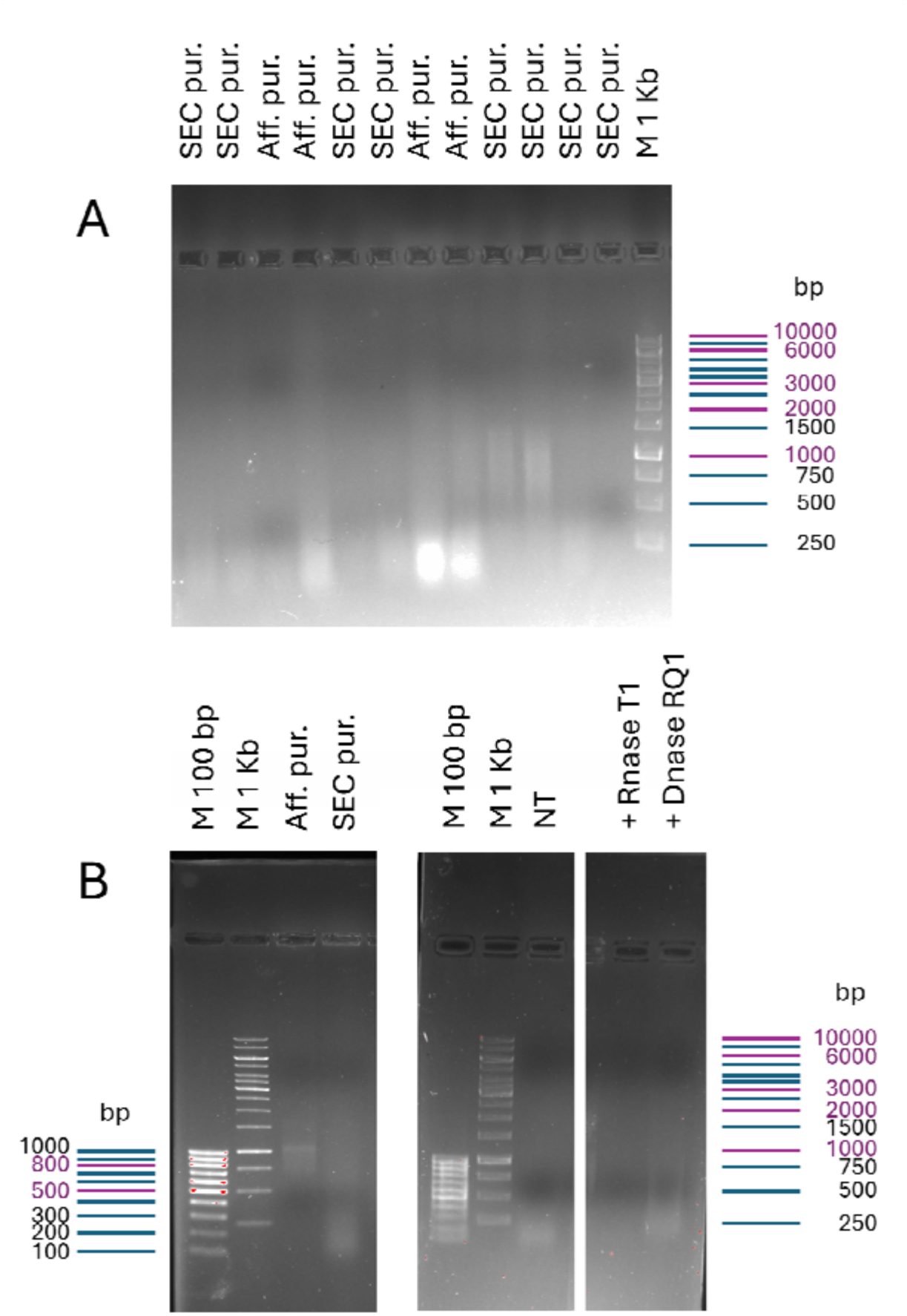
Detection and characterization of nucleic acids co-purified with GST-R4. **(A)** Representative samples collected at different stages of GST-R4 purification by GST-affinity chromatography and subsequent size-exclusion chromatography (SEC). Samples from selected fractions obtained during affinity purification and SEC were analyzed by agarose gel electrophoresis. The samples illustrate the persistence of nucleic acid material throughout the two purification steps. **(B)** Agarose gel electrophoresis of representative samples from GST-affinity purification (Aff. pur.) and SEC purification (SEC pur.), together with the SEC-purified sample subjected to enzymatic treatment. Aliquots (15 μL) of the SEC-purified sample were incubated for 30 min at 37°C either without enzyme (NT) or with RNase T1 or RQ1 DNase. The entire volume of each enzymatic reaction was loaded onto the gel. All samples were separated on 0.8% agarose gels in TAE buffer, stained with SYBR Safe and visualized using a Gel Doc imaging system. Molecular size markers (M) are indicated.

To characterize the nature of the co-purified nucleic acids, the most highly purified GST-R4 sample obtained after SEC was subjected to enzymatic treatment with RNase T1 or RQ1 DNase (Figure 4B). The untreated sample (NT) was analyzed in parallel. Treatment with RNase T1 markedly reduced the intensity of the nucleic acid band, whereas treatment with RQ1 DNase did not produce an appreciable change in the banding pattern compared with the untreated sample. This result supports the RNA nature of the co-purified nucleic acid material. This predominant nucleic acid band showed an apparent electrophoretic mobility corresponding approximately to □200 bp on a double-stranded DNA marker (Figure 4B). This apparent migration range is compatible with the presence of relatively long RNA species. All attempts to remove the associated RNA, by increasing ionic strength, adding divalent cations, or treating the protein preparation post-purification with nuclease, caused protein precipitation. These observations indicate that RNA removal is difficult under the conditions tested and suggest that RNA association contributes to protein stability. Based on the A260/A280 ratio and SEC stoichiometry, each GST-R4 molecule is estimated to engage □22–25 nt of RNA through its two zinc fingers, consistent with established TZF-RNA binding stoichiometries.

The thermal behavior of GST-R4 was further investigated under two ionic-strength conditions. The melting temperature of the protein was 56.5 ± 0.5°C at 200 mM NaCl, comparable to that of isolated GST (56°C), while at 1 M NaCl it decreased to 50.8 ± 0.5°C. This difference is consistent with the destabilization of the complex following RNA release at high ionic strength.

### 3.6 Functional monitoring of GST-R4 following affinity purification

The endoribonuclease activity of GST-R4 was assessed following GST-affinity purification as an additional functional quality-control step. Affinity-purified GST-R4 was incubated with a 5′-Cy5-labelled RNA substrate derived from the 3′UTR of human IL-6 mRNA, and substrate cleavage was monitored over a 120-min time course. While the substrate remained stable in cleavage buffer over time (Figure 5A), the GST-R4 preparation showed a progressive decrease in the full-length substrate signal, accompanied by the appearance of faster-migrating RNA species (Figure 5B). Additional bands with different electrophoretic mobility from the full-length substrate were also observed, highlighted the complexity of the sample. Therefore, size-exclusion chromatography (SEC) was subsequently introduced as a second purification step to further improve the purity of the GST-R4 preparation before its downstream biochemical and biophysical characterization.

**Figure 5.**
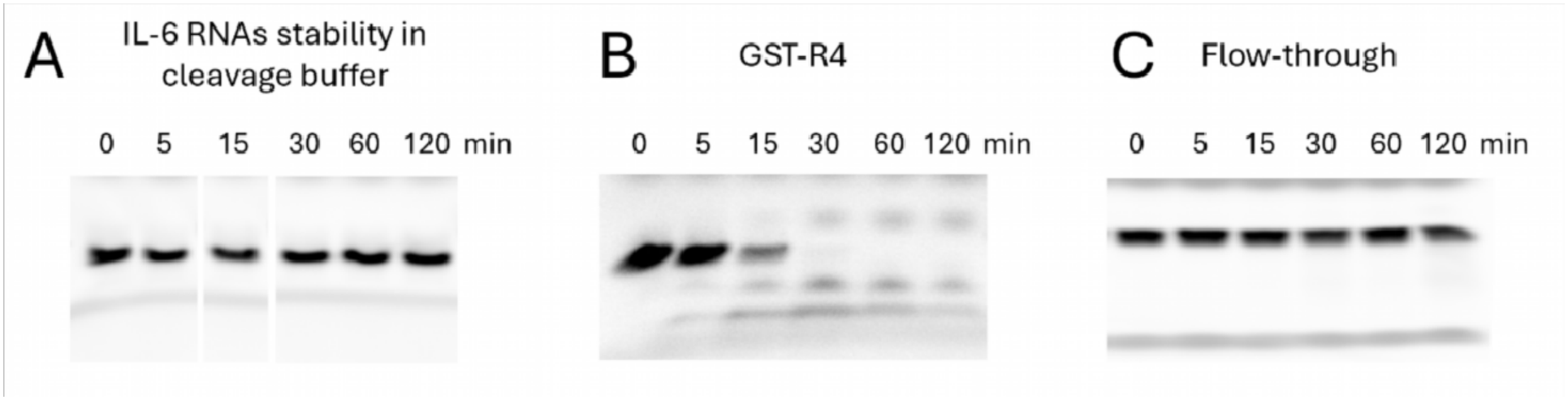
Time-course analysis of IL-6 RNA stability and cleavage by affinity-purified GST-R4. (A) Stability of the 5′-Cy5-labelled IL-6 RNA substrate incubated in cleavage buffer alone for the indicated time points (0–120 min). (B) Time-dependent changes in the electrophoretic profile of the IL-6 RNA substrate incubated with affinity-purified GST-R4. (C) Corresponding analysis performed with the flow-through material from the affinity purification. Reactions were incubated at 37°C, and aliquots were collected at the indicated time points. RNA substrates and products were separated by denaturing 12.8% polyacrylamide gel electrophoresis containing 7 M urea and visualized using a ChemiDoc imaging system.

To assess whether the observed RNA degradation could be attributed to contaminating bacterial RNases, control reactions were performed by adding an equivalent volume of flow-through material from the affinity purification to the RNA substrate (Figure 5C). IL-6 RNA remained substantially stable throughout the 120-min incubation, indicating that no detectable nonspecific RNase activity against this substrate was present in the flow-through under the conditions tested.

### 3.7 Crystallization trials

Screening of ~1000 crystallization conditions at protein concentrations of 2–6 mg/ml, using both sitting-drop and microbatch crystallization setup, did not yield protein crystals suitable for structure determination. The single candidate crystal (‘C9’), grown from a phosphate-based precipitant with 25% propanediol, pH 6.2 and 10% glycerol, exhibited unit-cell dimensions of a = 5.1 Å, b = 5.3 Å, c = 18.0 Å (orthorhombic) upon testing at ESRF beamline ID30A-3, indicating a small-molecule or salt crystal rather than a protein crystal. Additional trials incorporating sub-stoichiometric RNase A to reduce RNA-mediated heterogeneity also failed to produce diffracting crystals. The intrinsic conformational heterogeneity of the GST-R4/RNA complex, arising from the diversity of co-purified RNA sequences and their variable stoichiometry, is the most probable barrier to crystallization.

### 3.8 Preliminary cryo-EM analysis

Four cryo-EM grids prepared at GST-R4 concentrations of 0.5 and 1.0 mg/ml were screened on the FEI Talos Arctica 200 kV microscope. Representative micrographs show particles of approximately 10 nm diameter with morphologies consistent with folded macromolecular assemblies, along with elongated features attributable to RNA filaments (Figure 6).

**Figure 6.**
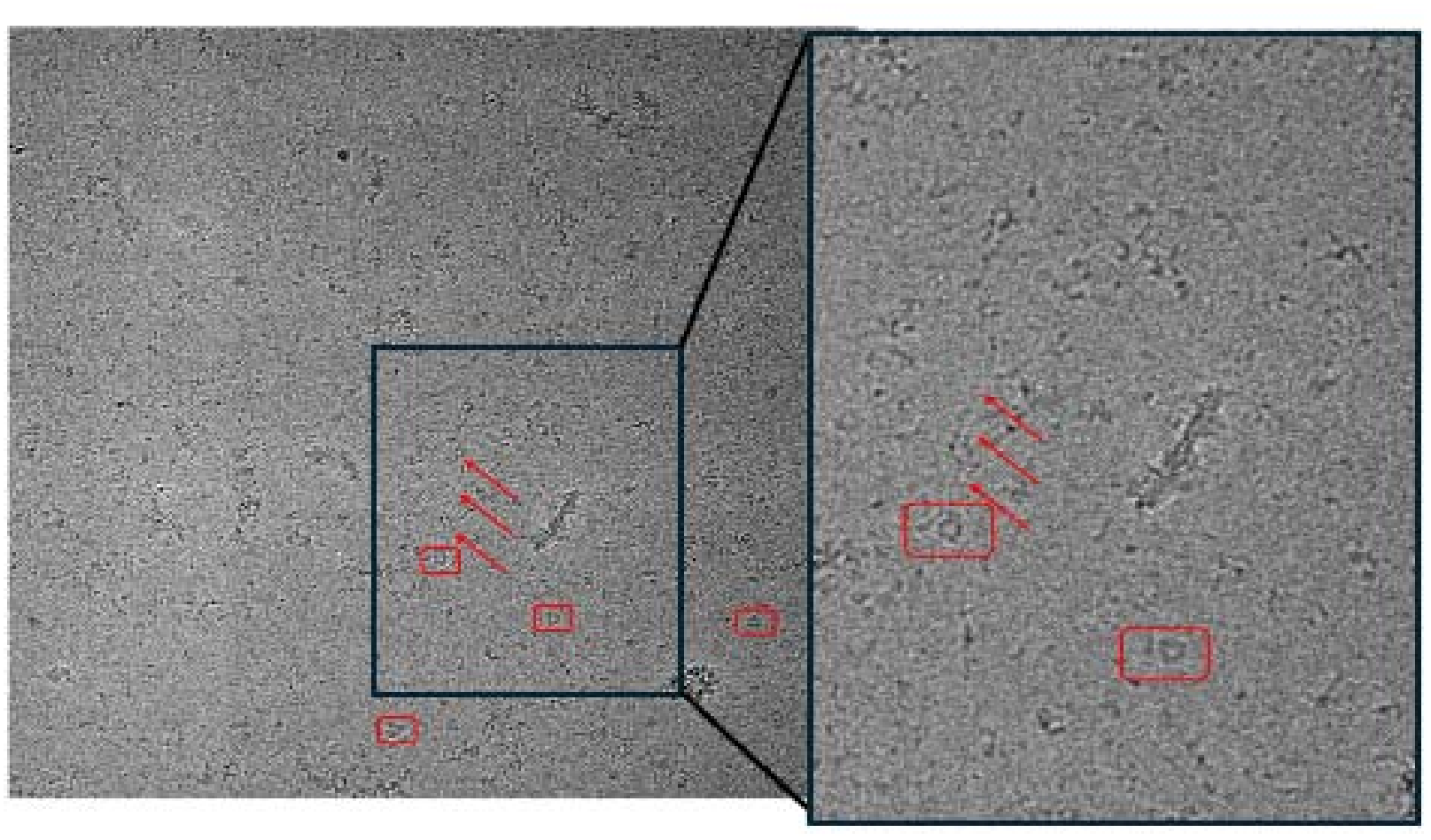
Representative electron micrograph of vitrified GST-R4 acquired using an FEI Talos Arctica transmission electron microscope operating at 200 kV. Red boxes highlight particles displaying a morphology consistent with a well-folded protein structure (estimated particle size: ~10 nm), while red arrows indicate structures that may correspond to RNA filaments. The right panel shows a magnified view of a selected region of the micrograph.

However, a substantial fraction of the sample appears partially denatured or aggregated, most likely as a result of air–water interface adsorption and mechanical stress during blotting and plunge-freezing. Even at the highest concentration tested (1.0 mg/ml), the density of apparently structured particles is insufficient for reliable 2D class averaging or 3D reconstruction. These observations identify vitrification optimization as the primary bottleneck for further structural work.

## 4. Discussion

The production of the engineered GST-R4 in soluble, active form required extensive optimization of construct design and expression conditions, and highlights challenges characteristic of multi-domain chimeric proteins. Among the five expression constructs tested, only the GST-fusion format proved productive. This strong dependence on the N-terminally GST fusion partner suggests that the tag acts not merely as an affinity handle but also as a co-translational solubility enhancer, potentially promoting the productive folding of the adjacent PIN domain. The requirement for low-temperature induction (16°C, overnight) further supports the notion that the chimeric architecture is inherently prone to aggregation during synthesis, a common feature of multi-domain proteins that require coordinated and sequential folding.

A major challenge encountered during GST-R4 production was its strong tendency to associate with endogenous bacterial RNA during expression. The sensitivity of the predominant nucleic acid species to RNase A, but not DNase I, identified it as RNA, with an estimated size of approximately 200 nt. This observation is compatible with an association between GST-R4 and bacterial transcripts during recombinant expression. This behavior is likely a direct consequence of its dual-domain architecture: the TZF domain retains RNA-binding activity and can capture endogenous bacterial transcripts in the reducing and crowded cytoplasmic environment, whereas the PIN domain does not efficiently cleave non-cognate RNA sequences. This RNA association is functionally informative, as it suggests that the zinc-finger motifs adopt a native-like conformation and retain their ability to engage RNA. Furthermore, co-purified RNA appears to play a structural stabilizing role, as attempts to remove the co-purified RNA were accompanied by protein precipitation. The ionic-strength-dependent decrease in melting temperature further highlights the sensitivity of GST-R4 thermal stability to the solution conditions. These properties impose important practical constraints on purification and sample handling: all purification steps must be completed rapidly at 4°C, RNA removal is not feasible without compromising protein stability, and GST-tag removal is hindered by the apparent inaccessibility of the protease cleavage sites within the protein–RNA complex.

A preliminary endoribonuclease assay performed after GST-affinity purification showed time-dependent changes in the electrophoretic profiles of the IL-6 RNA as substrate, indicating detectable RNA-cleavage activity in the GST-R4 preparation. Importantly, the corresponding flow-through controls showed no appreciable degradation of either substrate over the 120-min incubation, arguing against detectable nonspecific RNase contamination in the tested flow-through material under these conditions. Nevertheless, the additional RNA species observed in the affinity-purified preparations could not all be assigned unambiguously to cleavage products. These preliminary results therefore support the presence of detectable endoribonuclease activity in the affinity-purified GST-R4 preparation, while also highlighting the need for further purification and more detailed functional analysis.

Despite extensive optimization, GST-R4 could not be crystallized, even after screening a considerable number of crystallization conditions. This outcome is consistent with the conformational polydispersity likely introduced by heterogeneous co-purified RNA. The same heterogeneity may also account for the anomalously large apparent molecular weight (~570 kDa) observed by size-exclusion chromatography (SEC), suggesting that variable numbers of protein molecules assemble along RNA scaffolds of different lengths and sequences. Preliminary electron microscopy observations support this interpretation, revealing particles with an estimated size of approximately 10 nm, together with filamentous structures that may correspond to RNA. These observations are consistent with the SEC-based size estimates and provide initial morphological evidence of the protein–RNA assemblies formed by GST-R4.

Although preliminary cryo-EM analysis revealed promising features, current limitations, including low particle density on the grid and partial particle denaturation during vitrification, must be addressed before high-resolution structural studies can be undertaken. Potential improvements include optimization of grid surface treatments, such as graphene oxide or streptavidin-functionalized supports, to minimize surface adsorption, evaluation of alternative blotting protocols, and controlled enzymatic RNA digestion immediately before grid preparation. The latter approach may help reduce sample polydispersity while preserving the RNA-dependent structural stability of the protein complex.

Overall, this study establishes an optimized strategy for the recombinant expression and purification of GST-R4, an engineered chimeric PIN/TZF endoribonuclease. Among the five constructs and multiple expression conditions evaluated, only the GST-tagged R4 construct yielded appreciable amounts of soluble protein, with expression expressed in *E. coli* BL21(DE3) at 16°C providing the most suitable conditions. The optimized two-step purification strategy, combining GST-affinity chromatography and SEC within a single working day, produced protein with a purity exceeding 95%. Biochemical and biophysical analyses revealed the persistent co-purification of endogenous bacterial RNA and a large apparent hydrodynamic size. A preliminary assay following affinity purification also showed detectable endoribonuclease activity in the wild-type preparation. Although crystallization screening was unsuccessful, preliminary cryo-EM micrographs revealed folded particles of approximately 10 nm, providing a starting point for further structural investigations. Further optimization of sample homogeneity and vitrification conditions will be essential to obtain specimens suitable for high-resolution cryo-EM analysis. The expression, purification, and biochemical characterization strategies established in this work therefore provide a foundation for the next phase of structural studies on this novel class of chimeric enzymes.

## Acknowledgements

This work was supported by the Italian Ministry of Health (Bando Ricerca Finalizzata 2021) and by FISM (Fondazione Italiana Sclerosi Multipla) to LM and MM. The authors thank the cryo-EM facility of the Department of Biosciences, Università degli Studi di Milano, for access to the FEI Talos Arctica microscope. Synchrotron data collection was performed at the European Synchrotron Radiation Facility (ESRF), Grenoble, France, on beamline ID30A-3. EM also acknowledges support from PNRR Infrastructure (IR0000011, CUP B51E22000150006, “EBRAINS-Italy”), funded by MUR and the European Union – Next Generation EU, Mission 4, Component 2, Investment 1.3, for access to the protein production and characterization platform.

## CRediT Author Contribution Statement

Mario Milani: Conceptualization, Supervision, Writing original draft, Writing review & editing, Funding acquisition. Eloise Mastrangelo: Conceptualization, Supervision, Writing original draft, Writing review & editing, Funding acquisition. Paolo Cocomazzi: Investigation, Methodology, Writing review & editing. Silvia Ferrara: Resurses, Investigation, Methodology, Writing review & editing. Bianca Braggiotti: Investigation, Writing review & editing. Lucia Corcione: Investigation.

## Data Availability

The cryo-EM micrographs and synchrotron diffraction data reported in this article will be deposited in publicly accessible repositories upon acceptance. Raw cryo-EM micrographs will be deposited in the Electron Microscopy Public Image Archive (EMPIAR; https://www.ebi.ac.uk/empiar/). No three-dimensional reconstruction was achieved; therefore, no EMDB entry is applicable at this stage. X-ray diffraction images will be deposited in a public repository or made available upon reasonable request. All other data supporting the conclusions are presented within the article and its Supplementary Material.

## References

1. Akaki, K., Ogata, K., Yamauchi, Y., Iwai, N., Tse, K.M., Hia, F., Mochizuki, A., Ishihama, Y., Mino, T. & Takeuchi, O. (2021). IRAK1-dependent Regnase-1-14-3-3 complex formation controls Regnase-1-mediated mRNA decay. eLife, 10, e71966. DOI: 10.7554/eLife.71966.

2. Ferrara, S., Braggiotti, B., Mastrangelo, E., Di Gennaro, P., Bertoni, G. & Milani, M. (2025). Structural snapshots of the aldol condensation reaction of the enzyme trans-o-hydroxybenzylidenepyruvate hydratase-aldolase from Pseudomonas fluorescens N3. Biochemical and Biophysical Research Communications. DOI: 10.1016/j.bbrc.2024.151281.

3. Hudson, B.P., Martinez-Yamout, M.A., Dyson, H.J. & Wright, P.E. (2004). Recognition of the mRNA AU-rich element by the zinc finger domain of TIS11d. Nature Structural & Molecular Biology, 11(3), 257–264. DOI: 10.1038/nsmb738. PDB: 1RGO

4. Lai, W.S., Carballo, E., Strum, J.R., Kennington, E.A., Phillips, R.S. & Blackshear, P.J. (1999). Evidence that tristetraprolin binds to AU-rich elements and promotes the deadenylation and destabilization of tumor necrosis factor alpha mRNA. Molecular and Cellular Biology, 19(6), 4311–4323. DOI: 10.1128/MCB.19.6.4311.

5. Lindorff-Larsen, K., Piana, S., Palmo, K., Maragakis, P., Klepeis, J.L., Dror, R.O. & Shaw, D.E. (2010). Improved side-chain torsion potentials for the Amber ff99SB protein force field. Proteins: Structure, Function, and Bioinformatics, 78(8), 1950–1958. DOI: 10.1002/prot.22711.

6. Mahtani, K.R., Brook, M., Dean, J.L.E., Sully, G., Saklatvala, J. & Clark, A.R. (2001). Mitogen-activated protein kinase p38 controls the expression and posttranslational modification of tristetraprolin, a regulator of tumor necrosis factor alpha mRNA stability. Molecular and Cellular Biology, 21(19), 6461–6469. DOI: 10.1128/MCB.21.9.6461-6469.2001

7. Makita, S., Takatori, H. & Nakajima, H. (2021). Post-transcriptional Regulation of Immune Responses and Inflammatory Diseases by RNA-Binding ZFP36 Family Proteins. Frontiers in Immunology, 12, 711633. DOI: 10.3389/fimmu.2021.711633.

8. Mino, T., Murakawa, Y., Fukao, A., Vandenbon, A., Wessels, H.H., Ori, D., Uehata, T., Tartey, S., Akira, S., Suzuki, Y., Vinuesa, C.G., Ohler, U., Standley, D.M., Landthaler, M., Fujiwara, T. & Takeuchi, O. (2015). Regnase-1 and Roquin regulate a common element in inflammatory mRNAs by spatiotemporally distinct mechanisms. Cell, 161(5), 1058–1073. DOI: 10.1016/j.cell.2015.04.029. PDB: 5H9W.

9. Pronk, S., Páll, S., Schulz, R., Larsson, P., Bjelkmar, P., Apostolov, R., Shirts, M.R., Smith, J.C., Kasson, P.M., van der Spoel, D., Hess, B. & Lindahl, E. (2013). GROMACS 4.5: a high-throughput and highly parallel open source molecular simulation toolkit. Bioinformatics, 29(7), 845–854. DOI: 10.1093/bioinformatics/btt055.

10. Takeuchi, O. (2018). Endonuclease Regnase-1/Monocyte chemotactic protein-1-induced protein-1 (MCPIP1) in controlling immune responses and beyond. WIREs RNA, 9(1), e1449. DOI: 10.1002/wrna.1449.

11. Uehata, T. & Akira, S. (2013). mRNA degradation by the endoribonuclease Regnase-1/ZC3H12a/MCPIP-1. Biochimica et Biophysica Acta (BBA) - Gene Regulatory Mechanisms, 1829(6–7), 708–713. DOI: 10.1016/j.bbagrm.2013.03.001.

12. Xu, J., Peng, W., Sun, Y., Wang, X., Xu, Y., Li, X., Gao, G. & Rao, Z. (2012). Structural study of MCPIP1 N-terminal conserved domain reveals a PIN-like RNase. Nucleic Acids Research, 40(14), 6957–6965. DOI: 10.1093/nar/gks359. PDB: 3V33.

13. Yokogawa, M., Tsushima, T., Noda, N.N., Kumeta, H., Enokizono, Y., Yamashita, K., Standley, D.M., Takeuchi, O., Akira, S. & Inagaki, F. (2016). Structural basis for the regulation of enzymatic activity of Regnase-1 by domain-domain interactions. Scientific Reports, 6, 22324. DOI: 10.1038/srep22324.

